# Predicting chemotherapy response using a variational autoencoder approach

**DOI:** 10.1101/2021.01.04.425288

**Authors:** Qi Wei, Stephen A. Ramsey

## Abstract

**Motivation:** Multiple studies have shown the utility of transcriptome-wide RNA-seq profiles as features for machine learning-based prediction of response to chemotherapy in cancer. While tumor transcriptome profiles are publicly available for thousands of tumors for many cancer types, a relatively modest number of tumor profiles are clinically annotated for response to chemotherapy. The paucity of labeled examples and high dimension of the feature data limit performance for predicting therapeutic response using fully-supervised classification methods. Recently, multiple studies have established the utility of a deep neural network approach, the variational autoencoder (VAE), for generating meaningful latent features from original data. Here, we report first study of a semi-supervised approach using VAE-encoded tumor transcriptome features and regularized gradient boosted decision trees (XGBoost) to predict chemotherapy drug response for five cancer types: colon adenocarcinoma, pancreatic adenocarcinoma, bladder carcinoma, sarcoma, and breast invasive carcinoma.

**Results:** We found: (1) VAE-encoding of the tumor transcriptome preserves the cancer type identity of the tumor, suggesting preservation of biologically relevant information; and (2) as a feature-set for supervised classification to predict response-to-chemotherapy, the unsupervised VAE encoding of the tumor’s gene expression profile leads to better area under the receiver operating characteristic curve (AUROC) classification performance than either the original gene expression profile or the PCA principal components of the gene expression profile, in four out of five cancer types that we tested.

**Availability:** github.com/ATHED/VAE_for_chemotherapy_drug_response_prediction

**Supplementary information:** Supplementary data are available at *Bioinformatics* online.

## 1 Introduction

Although chemotherapy is a mainstay of treatment for aggressive cancers, many agents have serious side effects (Airley, 2009). Whether or not chemotherapy will provide a net benefit to a patient depends in large part on whether the malignancy responds to the treatment. Chemotherapy is often administered in cycles (Skeel, 2003), leading to multiple opportunities where treatment appropriateness may be (re-)assessed (Chabner and Longo, 2005). Currently, the medical cost-benefit of chemotherapy (versus a non-pharmaceutical approach) is assessed in light of patient health status, expected therapeutic tolerance, and tumor pathological classification (Kaestner and Sewell, 2007; Gurney, 2002).

For many cancer types, there is a broad spectrum of cases where the decision of whether or not to undergo or continue chemotherapy is difficult (Corrie, 2008; Whelan *et al.*, 2003; Malfuson *et al.*, 2008). The development of a quantitative model that could predict—based on a specific tumor’s molecular signature—whether or not the tumor will respond to chemotherapy would have significant clinical utility and would potentially improve patient quality-of-life. Moreover, an advance in machine-learning methods for the response-to-chemotherapy prediction problem (Chiu *et al.*, 2019; Geeleher *et al.*, 2014) would have potential crossover benefits for other prediction problems in precision medicine.

Oncogenesis is driven by alterations in the somatic genome and epigenome in cancer cells (Weir *et al.*, 2004); however, the somatic genetic or epigenetic determinants of response to chemotherapy are also thought to exert measurable effects on gene expression in the tumor. Consistent with this theory, studies of various cancer types have demonstrated that biomarkers identified from systematic measurement of the patient’s cancer transcriptome or proteome correlate with the probability that a tumor will respond to chemotherapy, for example, a five-protein signature in breast cancer (Gámez-Pozo *et al.*, 2017), 13- and 14-gene signatures in rectal cancer (Casado *et al.*, 2011; Del Rio *et al.*, 2007), and a 63-gene signature in liver cancer (Kurokawa *et al.*, 2004). Taken together, the findings from such “omics” biomarker studies suggest that RNA sequencing-(RNA-seq (Wang *et al.*, 2009))-based transcriptome measurements of tumor samples labeled with clinical response can be used to train machine-learning classifiers for predicting response to chemotherapy. However, the accuracy of such models is presently limited by the small number of available training cases that are labeled for clinical outcome, given the large size of the transcriptome (∼60k genes Frankish *et al.*, 2018) and the significant intertumoral variance of gene expression. For typical cancers, most of the profiled tumor transcriptomes are not labeled for chemotherapeutic response; the ratio of such unlabeled to labeled tumor datasets in the Cancer Genome Atlas (TCGA) dataset (Hutter and Zenklusen, 2018) ranges from 10–20, depending on the cancer type. While using (exclusively) supervised learning methods for the response-to-chemotherapy prediction problem has been a sensible first step, *unlabeled data* are a substantial resource that could—in the context of a *semi-supervised* approach—reveal multivariate structure or patterns that could ultimately improve predictive accuracy. Semi-supervised approaches that fuse unsupervised data reduction methods (such as principal components analysis, or PCA) for low-dimensional embedding with supervised methods (such as decision trees) for prediction have proved beneficial in problems where large unlabeled datasets are available, for example, a PCA-XGBoost method has been previously used in finance (Wen and Huang, 2020), and an independent components analysis-based method has been used to classify electroencephalographic signals (Qin *et al.*, 2006).

Multiple studies (An and Cho, 2015; Li and She, 2017; Bouchacourt *et al.*, 2017; Kipf and Welling, 2016) have established the power of the variational autoencoder (VAE; Kingma and Welling (2013); Jimenez Rezende *et al.* (2014))—an unsupervised nonlinear data embedding model with two deep neural networks oppositely connected through a low-dimensional probabilistic latent space—for finding meaningful and useful latent features in high-dimensional data. In the context of cancer bioinformatics, VAEs have been variously used to (i) model cancer gene expression and capture biologically-relevant features using the TCGA Pan-cancer Project RNA-seq dataset (Way and Greene, 2018); (ii) find encodings that correlate with biological features such as patient sex and tumor type (Titus *et al.*, 2018); (iii) find encodings that can be used to predict gene inactivation in cancer (Way and Greene, 2017); and (iv) obtain an encoding that is predictive of chemotherapy resistance (George and Lio, 2019). Based on their exploration of multiple VAE architectures for predicting gene inactivation in a pan-cancer dataset, Way & Greene reported (2017) biological insights obtained from the latent-space embeddings learned by VAEs. George and Lio (2019) used a VAE-based, fully unsupervised approach to encode ovarian tumor transcriptomes and obtained latent-space features that were associated with response to chemotherapy. These studies suggest that a tumor transcriptome VAE may be broadly useful for the response-to-chemotherapy prediction problem and they set the stage for the present multi-cancer investigation of the utility of the tumor transcriptome VAE in precision oncology.

Given previous reports of success using a VAE to obtain useful low-dimensional encodings of transcriptome data (Dong *et al.*, 2020; Way and Greene, 2018; Way and Greene, 2017), in this work, we first sought to ascertain to what extent a VAE encoding of tumor transcriptome data would preserve biological characteristics—spanning multiple genes at a time that have coordinated variation across tumors— that are associated with distinct cancer types. To answer this question, we trained a pan-cancer transcriptome VAE and used it to encode TCGA tumor RNA-seq data from 9,310 tumors comprising 32 different cancer types, focusing on the top 5,000 most variable genes. We trained the VAE using an efficient contemporary optimization engine (Adam) to find the VAE coefficient values that together balance reconstruction loss and desired latent-space distributional shape. We applied an unsupervised two-dimensional embedding method (*t*-distributed stochastic neighbor embedding, or *t*-SNE) directly to tumor transcriptome and to the VAE-embedded tumor transcriptome data, and mapped clusters of tumors by cancer type across the two *t*-SNE embeddings. We found (Sec. 2.1) that the VAE preserves the clustering of tumors of the same cancer type, suggesting biological fidelity in the components of the VAE embedding.

Next, to set the stage for a semi-supervised approach for predicting cancer response to chemotherapy, we selected five cancer types (breast, bladder, colon, pancreatic, and sarcoma) based on sufficient availability of clinically labeled data and then defined three different VAE architectures: `VAE-1`, which we used to obtain feature data for bladder, breast, and pancreatic cancer; `VAE-2`, for sarcoma; and `VAE-3`, for colon cancer. In order to train a VAE, it is necessary to specify a reconstruction loss function; both L2 and L1 reconstruction loss have been used for training VAEs in machine-learning, and we sought to clarify which is best for this application. Thus, we trained each of the three VAE architectures on 2,606 tumor transcriptomes from TCGA, in an unsupervised fashion, separately using L1 loss and L2 loss. Next, in order to label tumors for response to chemotherapy, we analyzed the available TCGA clinical data regarding the outcome of pharmaceutical therapy (in most cases including chemotherapy) for each of the patients, and thereby assigned a label “responded” or “progressive” to 806 out of the 2,606 tumors (Sec. 2.2); the remainder of the tumors were unlabeled and thus used only during VAE training. For the 806 labeled tumors, we used the VAE-encoded latent vectors as feature data for supervised prediction of the binary label using gradient boosted decision trees (XGBoost; Chen and Guestrin (2016)). Using this semi-supervised “VAE-XGBoost” approach, we found (Sec. 2.3) that a VAE trained using L1 reconstruction loss yields features that result in better classification performance (by area under the receiver operating characteristic, AUROC) than a VAE trained using L2.

In the main part of this work, using XGBoost, we measured response-to-chemotherapy prediction performance for each of three tumor transcriptome-derived feature sets: (i) expression levels of the top 20% of genes, by intertumoral variance (a fully supervised approach); (ii) the first 387 principal components of expression levels of “top 20%” genes (“semi-supervised PCA-XGBoost”); and (iii) VAE-encoded expression levels of the top 20% genes (“semi-supervised VAE-XGBoost”, our new method, Fig. 1). Within a cross-validation framework for AUROC performance evaluation, we found (Sec. 2.4) that for four out of five cancer types, the semi-supervised VAE-XGBoost approach outperformed the fully-supervised approach. Moreover, for four out of the five cancer types, semi-supervised VAE-XGBoost outperformed semi-supervised PCA-XGBoost. Finally, for the one cancer type for which PCA-XGBoost outperformed VAE-XGBoost, we investigated their relative performance through the lens of XGBoost feature importance (Sec. 2.5). Below, we describe our results (Sec. 2) and the VAE-XGBoost method in detail (Sec. 5).

**Fig. 1:**
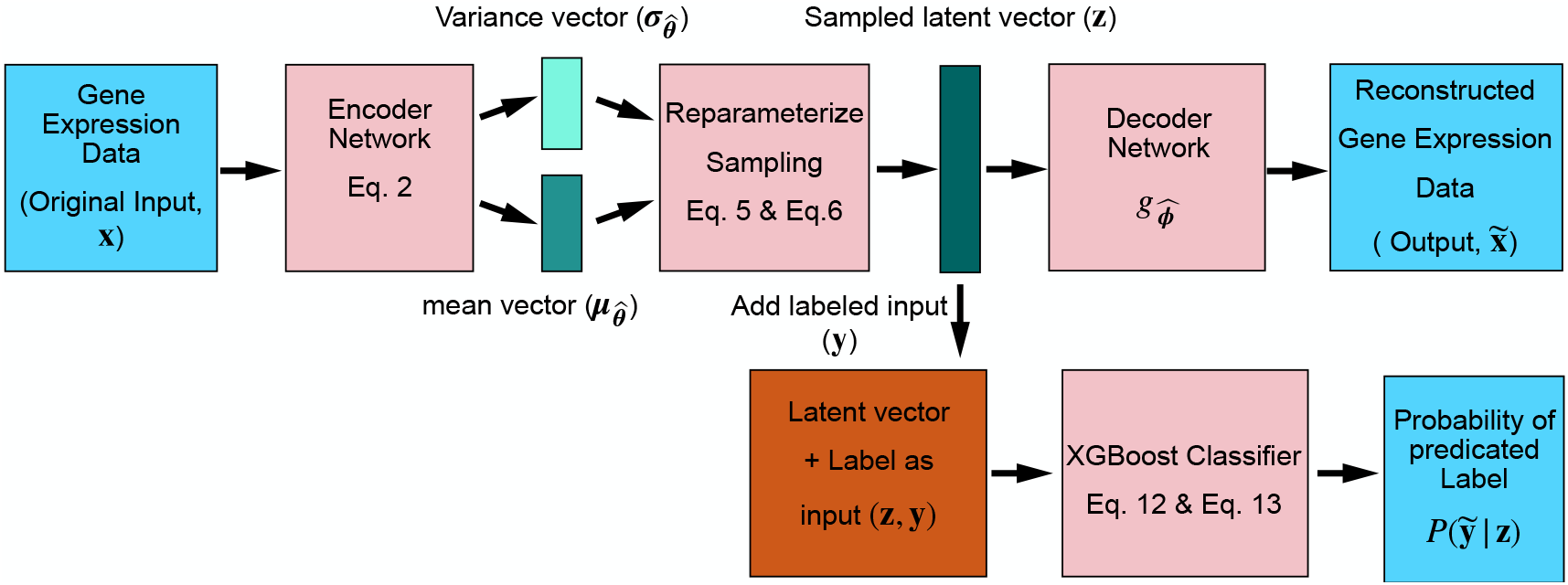
Overview of the VAE-XGBoost method that we used for predicting tumor response to chemotherapy. For each tumor *t*, the encoder’s input vector ***x***_*t*_ contains the levels of the top 20% of genes by intertumoral gene expression variance (Sec. 5.1). Each network has multiple fully connected dense layers (Sec. 5.5). The encoder outputs two vectors of configurable latent variable dimension *h* ≪ *m* (Sec. 5.5): a vector of means ***μ*** and a vector of standard deviations ***σ*** that parameterize the multivariate normal latent-space vector ***Z***|***x***_*t*_ (Sec. 5.3). The sampled encoding ***Z***|***x***_*t*_ = ***z***_*t*_ is passed to the decoding neural network (decoder), whose architecture is identical to (with inversion) that of the encoder network. The sampled latent-space vector ***z*** *_t_* is passed to XGBoost for supervised classification to predict response to chemotherapy (training label ***y***, prediction 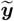).

## 2 Results

### 2.1 VAE encoding preserves cancer type features

Given multiple reports (Dolezal *et al.*, 2018; Esteva *et al.*, 2017) that *t*-SNE can be used to visualize the grouping of cancer types from high-dimensional molecular tumor data, we investigated the extent to which

VAE encoding of tumor transcriptomes preserves data-space features that determine cancer type-specific groupings. In order to do so, we obtained (Sec. 5.1) from the TCGA data portal RNA-seq transcriptome data for 9,310 tumors labeled for 32 different cancer types (listed in Fig. 2). As a baseline view of transcriptome-based cancer type groupings, we generated a two-dimensional embedding of the 9,310 tumor samples by applying *t*-SNE (Sec. 5.2) to the expression levels of the top 5,000 most variable genes, yielding 32 distinct clusters (Fig. 2A). Next, we trained (Sec. 5.3) a VAE to encode the expression levels of the 5,000 most variable genes in each of 9,310 tumors into 9,310 points in a 50-dimensional latent space. An unsupervised *t*-SNE visualization (Fig 2B) of the VAE-encoded tumor transcriptome data was remarkably similar in structure to the *t*-SNE visualization of the 5,000-dimensional original dataset, with intercluster distances for all pairs of clusters correlated between of the two *t*-SNE plots (*R* = 0.49; see Fig. S1). This analysis indicated that the VAE encoding preserves data-space features that distinguish individual cancer types.

**Fig. 2:**
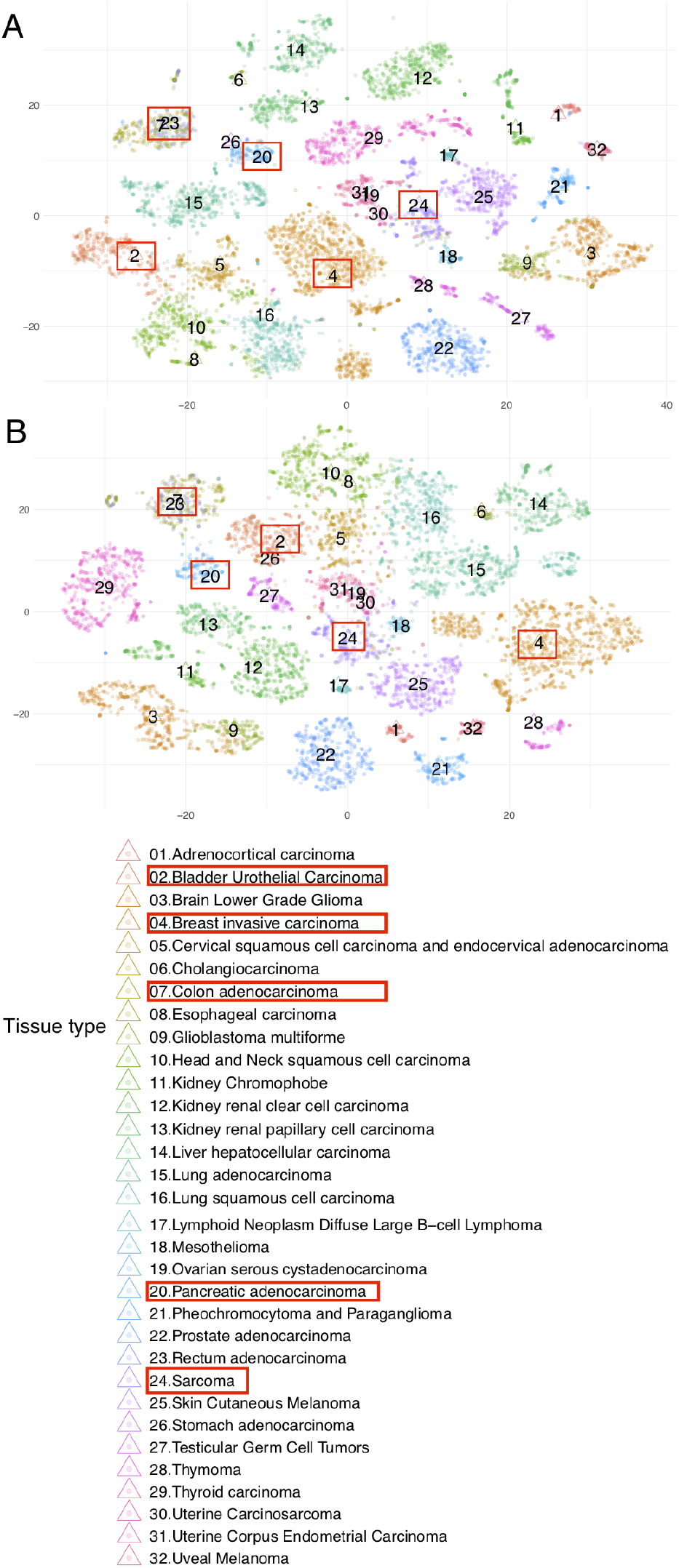
Marks represent tumor transcriptomes visualized using *t*-SNE, with colors representing cancer types. (A) Original gene expression data of the top 5,000 most variable genes. (B) VAE compressed gene expression data. Red rectangles denote the five cancer types selected for chemotherapy response classification (Sec. 2.4).

### 2.2 Obtaining a labeled tumor transcriptome dataset

Having demonstrated that the VAE can efficiently encode tumor transcriptomes while preserving features that distinguish different cancer types, and to set the stage for implementing a semi-supervised approach for predicting response to chemotherapy, we obtained a five-cancer-type tumor transcriptome dataset with a significant subset of the tumors labeled for “response to chemotherapy”, as described below. We obtained transcriptomes of 806 tumors across five cancer types [colon adenocarcinoma (COAD), pancreatic adenocarcinoma (PAAD), bladder carcinoma (BLCA), sarcoma (SARC), and breast invasive carcinoma (BRCA); see Table 1] that we selected based on availability of a sufficient amount of labeled data in TCGA (see Sec. 5.1) and generated binary clinical labels for them corresponding to “responded” or “progressive” (see Sec. 5.4). Among these tumors, the class balance ratio, i.e., the ratio of responding tumors to progressive disease tumors, ranged from a low of 0.77 for pancreatic cancer to a high of 8.61 for breast cancer.

**Table 1.**
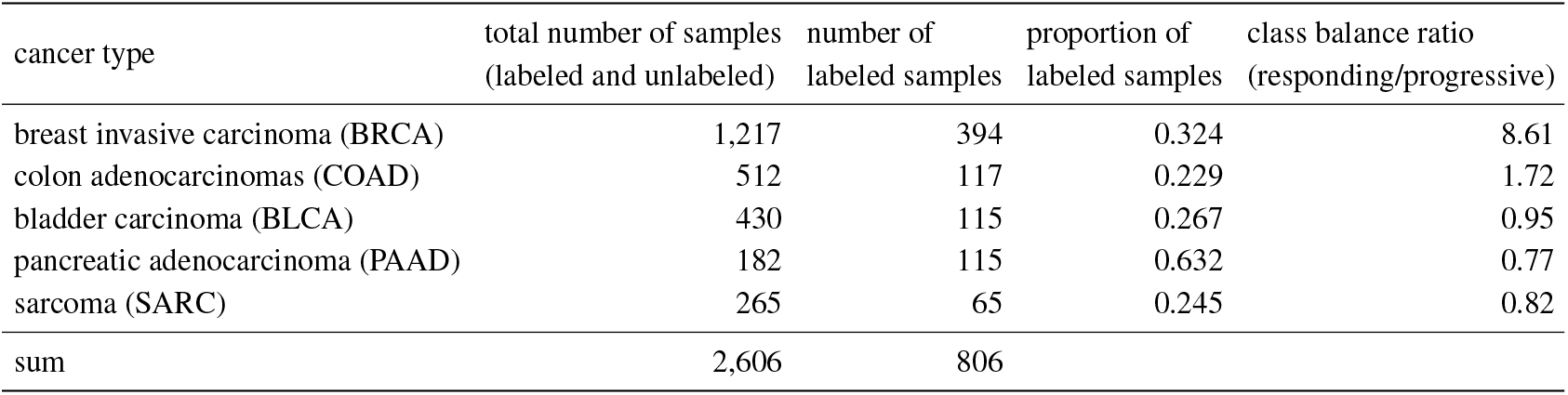
Table of numbers of samples with chemotherapy response record for each cancer type (n.b., the total number of labeled tumor samples exceeds the total number of patients because some patients had multiple tumors). After each cancer type, its TCGA abbreviation is shown in parentheses.

| cancer type | total number of samples<br>(labeled and unlabeled) | number of<br>labeled samples | proportion of<br>labeled samples | class balance ratio<br>(responding/progressive) |
| --- | --- | --- | --- | --- |
| breast invasive carcinoma (BRCA) | 1,217 | 394 | 0.324 | 8.61 |
| colon adenocarcinomas (COAD) | 512 | 117 | 0.229 | 1.72 |
| bladder carcinoma (BLCA) | 430 | 115 | 0.267 | 0.95 |
| pancreatic adenocarcinoma (PAAD) | 182 | 115 | 0.632 | 0.77 |
| sarcoma (SARC) | 265 | 65 | 0.245 | 0.82 |
| sum | 2,606 | 806 |  |  |

### 2.3 L1 loss is better than L2 loss for this application

Having obtained 2,606 tumor transcriptomes across five cancer types with 806 of the tumors labeled for response to chemotherapy, we next sought to determine which type of VAE reconstruction loss function—L1 loss or L2 loss—would yield transcriptome encodings that are most amenable to accurate XGBoost-based prediction of response to chemotherapy. On the 2,606 tumor transcriptomes, we trained two sets of cancer type-specific VAEs (see Sec. 5.5) using L1 and L2 loss functions, respectively. We used the L1 and L2 VAEs to encode the 806 labeled tumor transcriptomes (the top 20% most variable genes in each cancer type, merged across the five cancers, for a total of 13,584 genes) spanning the five cancer types, yielding (for each cancer type) two feature matrices (one for L1 loss and one for L2 loss) that we separately evaluated for XGBoost prediction (Sec. 5.6) of the binary response-to-chemotherapy class label. By test-set area under the receiver operating characteristic (AUROC; Sec. 5.7), averaged across the five cancers, we found (Fig. 3) that the features that were generated by the L1 VAEs led to 6.2% better (*p* < 10^−9^, Welch’s *t*-test) classification performance than the features generated by the L2 VAEs, and thus, for all subsequent analyses, we used VAEs trained with L1 loss.

**Fig. 3:**
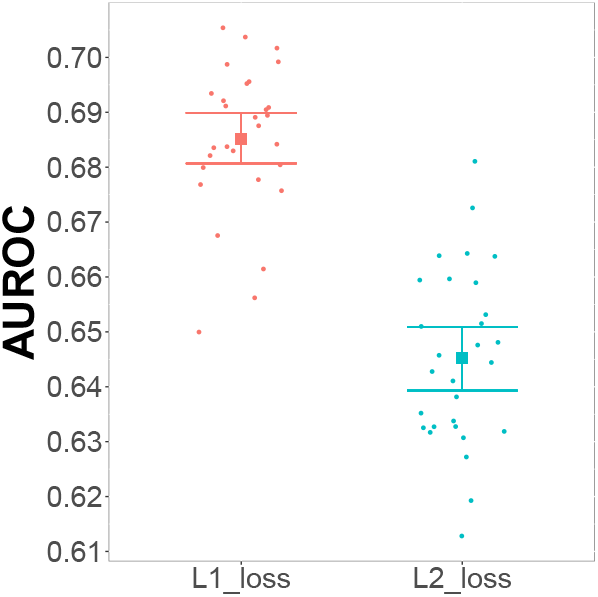
Average AUROC results over five different types of cancer, by loss type. Squares, mean values; bars, 95% confidence interval (c.i.).

### 2.4 Chemotherapy drug response classification result

Having selected L1 reconstruction loss for training VAEs to encode tumor transcriptomes for predicting response-to-chemotherapy, we focused on the key question of whether (and to what extent) a semi-supervised approach using the VAE can outperform (in terms of predictive accuracy) a fully supervised approach or a semi-supervised approach based on a traditional dimensional reduction technique (principal components analysis, PCA). In brief, our VAE-based semi-supervised approach involves three steps: (i) training a VAE to encode clinically *unlabeled* tumor transcriptomes (for the top 20% most variable genes) for a single cancer type, into a low-dimensional space (Sec. 5.5); (ii) using that VAE to obtain latent-space encodings for the tumor transcriptomes that are labeled for a relevant clinical endpoint (in this work, response to chemotherapy); and (iii) training and testing a supervised classifier (in this work, XGBoost binary classification) using the latent-space encodings as feature data. To address the question of whether this VAE-based, semi-supervised (VAE-XGBoost) approach can outperform a fully supervised approach, we compared the performance of the VAE-XGBoost method to a fully supervised approach in which we applied XGBoost directly to the tumor expression levels of the top 20% most variable genes (13,584 genes) as feature data. In the same analysis, to address the question of whether the VAE-XGBoost method could outperform a semi-supervised approach based on PCA dimensional reduction, we compared the VAE-XGBoost method to the PCA-XGBoost method. We carried out this analysis for each of the five cancer types separately, using the set of cancer type-specific labeled tumors (totaling 806 labeled tumors). We measured performance using test-set AUROC in a cross-validation framework (Sec. 5.7).

For four out of five cancer types (breast, colon, pancreatic, and sarcoma), in terms of test-set AUROC, the VAE-XGBoost approach outperformed the fully-supervised approach of applying XGBoost directly to the expression levels of the tumors’ top 20% most variable genes (Fig. 4), by both Welch’s *t*-test and Wilcoxon’s signed-rank test (Table 2); for BLCA, the semi-supervised VAE-XGBoost and fully-supervised models’ performances were statistically indistinguishable. Additionally, for four out of five cancer types (bladder, breast, pancreatic, and sarcoma), the semi-supervised VAE-XGBoost method significantly outperformed the semi-supervised PCA-XGBoost method (Fig. 4 and Table 2). The five-cancer average AUROC for VAE-XGBoost was 0.682, a performance gain of 5.4% over the five-cancer average AUROC for PCA-XGBoost (0.646) and a gain of 3.6% over the fully-supervised model’s average (0.658). Notably, a single deep VAE architecture (`VAE-1`, which had a 50-dimensional latent space and six layers in the encoder; see Sec. 5.5) yielded latent-space encodings that outperformed semi-supervised PCA-XGBoost for three cancer types (bladder, breast, and pancreatic).

**Table 2.**
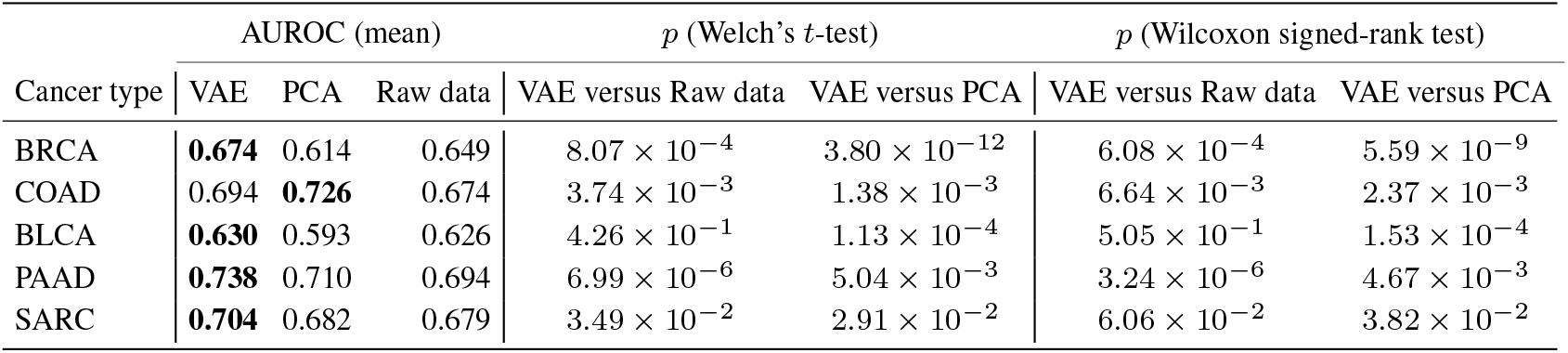
Quantitative AUROC performances of XGBoost (“Raw data”), PCA-XGBoost (“PCA”), and VAE-XGBoost (“VAE”), along with pairwise comparisons.

| Cancer type | AUROC (mean) | | | $p$ (Welch’s $t$-test) | | $p$ (Wilcoxon signed-rank test) | |
| --- | --- | --- | --- | --- | --- | --- | --- |
|  | VAE | PCA | Raw data | VAE versus Raw data | VAE versus PCA | VAE versus Raw data | VAE versus PCA |
| BRCA | **0.674** | 0.614 | 0.649 | $8.07 \times 10^{-4}$ | $3.80 \times 10^{-12}$ | $6.08 \times 10^{-4}$ | $5.59 \times 10^{-9}$ |
| COAD | 0.694 | **0.726** | 0.674 | $3.74 \times 10^{-3}$ | $1.38 \times 10^{-3}$ | $6.64 \times 10^{-3}$ | $2.37 \times 10^{-3}$ |
| BLCA | **0.630** | 0.593 | 0.626 | $4.26 \times 10^{-1}$ | $1.13 \times 10^{-4}$ | $5.05 \times 10^{-1}$ | $1.53 \times 10^{-4}$ |
| PAAD | **0.738** | 0.710 | 0.694 | $6.99 \times 10^{-6}$ | $5.04 \times 10^{-3}$ | $3.24 \times 10^{-6}$ | $4.67 \times 10^{-3}$ |
| SARC | **0.704** | 0.682 | 0.679 | $3.49 \times 10^{-2}$ | $2.91 \times 10^{-2}$ | $6.06 \times 10^{-2}$ | $3.82 \times 10^{-2}$ |

**Fig. 4:**
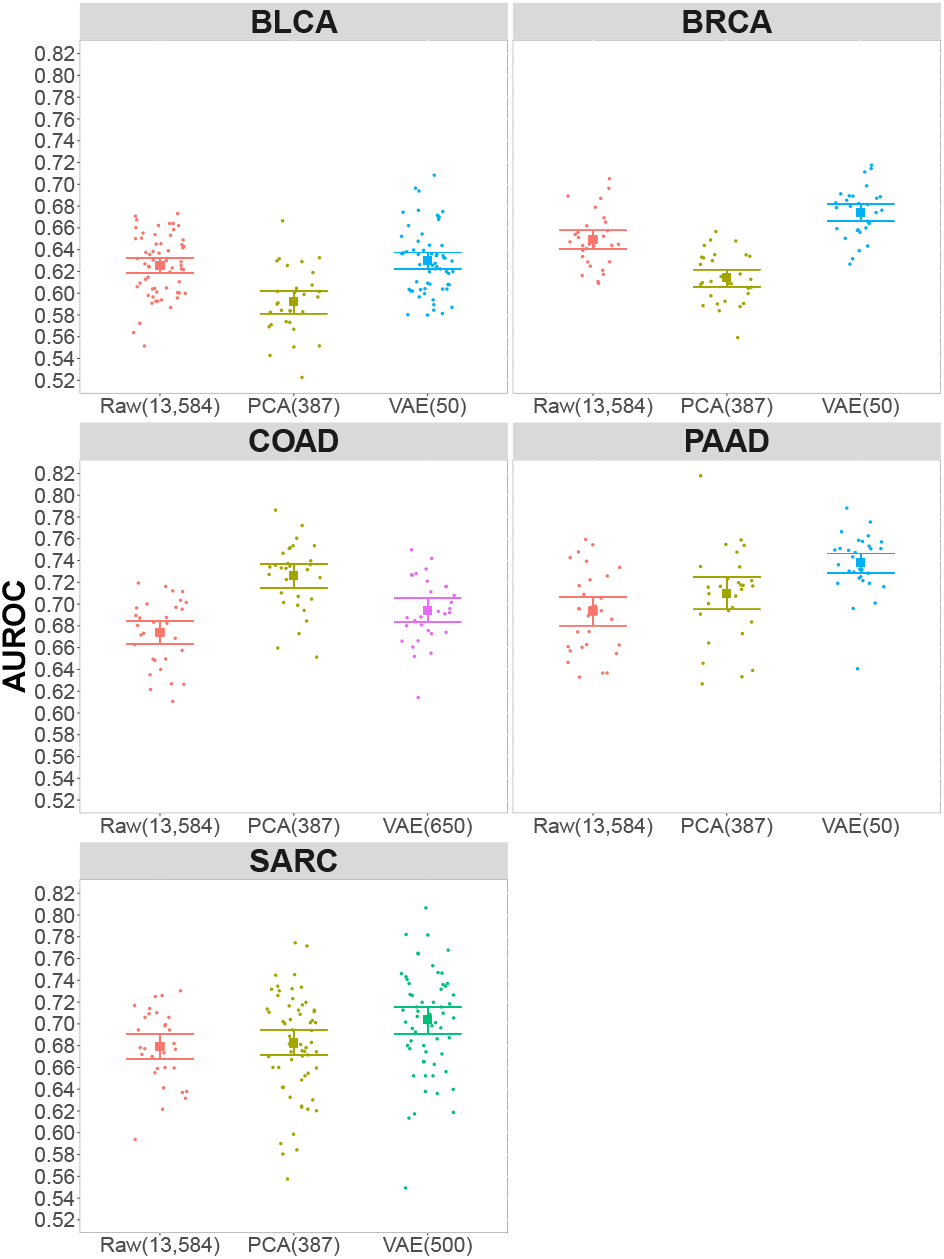
Test-set performance of the three models for predicting response to chemotherapy, across five cancer types. Group abbreviations: “PCA(387)”, the PCA-XGBoost semi-supervised method (387: number of principal components used as features); “Raw(13,584)“, the fully-supervised XGBoost method (13,584: number of genes used as features); and “VAE(*n*)”, the VAE-XGBoost semi-supervised method (*n*: dimension of the latent feature space). Marks correspond to individual replications of five-fold cross-validation; solid squares denote mean; bars indicate 95% c.i; colors denote the type of feature-set (Sec. 5.5): red, “PCA”; olive, “Raw”; cyan, `VAE-1`; magenta, `VAE-2`; green, `VAE-3`.

### 2.5 PCA & VAE feature importance scores, for COAD

Having established that the semi-supervised VAE-XGBoost outperforms the semi-supervised PCA-XGBoost approach for tumor transcriptome-based prediction of response to chemotherapy for four out of five cancer types, we sought to understand the basis for the higher performance of PCA-XGBoost over VAE-XGBoost on the fifth cancer type, colon adenocarcinoma (COAD). Specifically, we investigated whether the strong performance of PCA-XGBoost on COAD is attributable to differences in the distributions of XGBoost feature importance scores (Sec. 5.6) of the PCA features versus VAE latent-space features. We found that the distribution of feature importance scores (as a function of rank) was more sharply peaked at lowest-ranked features in the VAE than in the PCA (Fig. 5), suggesting that the performance gain with PCA reflects a broader spectrum of informative features for that particular cancer type.

**Fig. 5:**
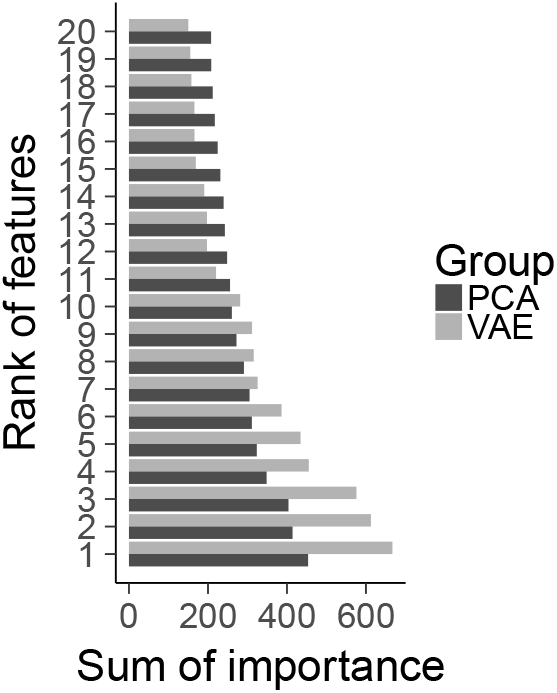
Bars indicate the sum (over 30 replications) of XGBoost feature importance scores. “Group” indicates the low-dimensional embedding method used (VAE or PCA). Bars separately ordered from highest to lowest (only top 20 most important features shown).

## 3 Discussion

As far as we are aware, this work is the first report of a broad (five-cancer) investigation of the potential for a VAE-based, semi-supervised approach for predicting response to chemotherapy. Across the five cancer types that we studied, the ratio of responding tumors to progressive disease tumors ranged from a low of 0.77 for pancreatic cancer to a high of 8.61 for breast cancer, reflecting a broad range of resistances to standard-of-care chemotherapy. Our results clearly demonstrate the utility of the VAE for compressing high-dimensional data to a continuous, low-dimensional latent space while retaining features that are essential for distinguishing different cancer types and for predicting response to chemotherapy. Nevertheless, three limitations of this work bear noting.

The first limitation concerns the type(s) of tumor “omics” data from which features are derived for the predictive model. While in this work we focused on tumor transcriptome data which can be measured with high precision over a wide dynamic range of transcript abundances by RNA-seq, we note that TCGA datasets of tumor somatic mutations and copy number alteration events are also available (Hutter and Zenklusen, 2018). Given the voluminous literature on the use of tumor somatic genomic data for precision cancer diagnosis (Mitchel *et al.*, 2019; Zhang *et al.*, 2020; Lee *et al.*, 2019), tumor DNA datasets are fertile ground for developing a semi-supervised, multi-omics model for predicting response to chemotherapy.

Second, we noted for decision tree-based response-to-chemotherapy prediction, the performance of VAE-encoded transcriptome features is somewhat sensitive to the type of normalization used for the input data (data not shown). We explored various types of normalization for the RNA-seq data including standardization of log counts and using FPKM data, we ultimately chose min-max-normalized log_2_ total-count-normalized counts (Sec. 5.1) for the gene expression levels to be used to derive features. However, there are additional transcript quantification methods (Evans *et al.*, 2017) that could be explored in the context of finding optimal tumor transcriptome VAE encodings for precision oncology. A similar comment applies to the specific form of the reconstruction loss function: in our analysis, features from the VAE trained with L1 loss clearly (across five cancers) outperformed those from the VAE trained with L2 loss, and thus, consistent with Way and Greene (2017), we used L1 loss for the VAE that we used to address the main question of this work (Sec. 2.4) as well as the pan-cancer *t*-SNE analysis (Sec. 2.1)

The third limitation relates to the VAE architecture. While it is promising that a single deep VAE architecture (VAE-1, with a 50-dimensional latent space and six fully-connected layers) yielded features that outperformend PCA and the original RNA-seq feature data for three different cancer types (bladder, breast, and pancreatic), for colon cancer and sarcoma, it was necessary to use shallower (two-layer) VAE architectures with bigger latent space dimensions (650 and 500, respectively). Of the five cancers studied, colon cancer and sarcoma had the lowest proportions of labeled samples (0.229 and 0.245 respectively; see Table 1). Our findings suggest that for some cancers, a deep, low-latent-dimension VAE architecture yields optimal features for predicting response, while for other cancers, a shallow, medium-sized-latent-dimension VAE architecture is more effective. More study with larger datasets will be required in order to determine whether a single VAE architecture could be successfully used for general-purpose tumor transcriptome feature extraction for precision oncology.

While our results show promise for the VAE in the context of a semi-supervised approach for response-to-chemotherapy prediction, for colon cancer, the VAE-XGBoost method did not outperform PCA-XGBoost (though it did outperform the fully supervised approach of XGBoost trained on the unencoded gene expression data). One possible explanation for the colon cancer-specific superior performance of PCA features over VAE features for predicting response to chemotherapy may be due to the fact that while (for COAD) feature importance for the VAE features is sharply peaked for the first few features and falls off fairly rapidly with feature rank, the PCA features have a much flatter distribution of relative feature importance (Fig. 5). Follow-on studies with larger datasets will be required to delineate under what circumstances transcriptome VAE encodings will prove superior to linear principal components.

## 4 Conclusions

For four of the five cancer types that we studied, the semi-supervised VAE-XGBoost approach significantly outperformed a semi-supervised PCA-XGBoost approach for tumor transcriptome-based prediction of response to chemotherapy, reaching a top AUROC of 0.738 for pancreatic adenocarcinoma. For four out of five cancer types, the semi-supervised VAE-XGBoost approach significantly outperformed a fully-supervised approach consisting of XGBoost applied to the expression levels of the top 20% most variably expressed genes. Given high-dimensional “omics” data, the VAE is a powerful tool for obtaining a nonlinear low-dimensional embedding; it yields features that retain biological patterns that distinguish between different types of cancer and that enable more accurate tumor transcriptome-based prediction of response to chemotherapy than would be possible using the original data or their principal components.

## 5 Methods

We carried out all data processing and machine-learning tasks on a Dell XPS 8700 workstation equipped with Nvidia Titan RTX GPU and running the Ubuntu GNU/Linux operating system version 16.04. All of the analysis code that we implemented was executed in Python version 3.5.5 except that we used R version 3.3.3 for statistical analysis of AUROC values (Sec. 5.7), gene-level MAD calculations (Sec. 5.1) and plotting (Sec. 5.2). We carried out all statistical tests using the R computing environment (version 3.3.3) and using the R software package `stats` version 3.4.4.

### 5.1 Gene expression data

From the Xena data portal (Goldman *et al.*, 2019), we obtained TCGA Level 3 tumor RNA-seq transcriptome data of 32 cancer types (totaling 9, 310 tumors) and, for the response-to-chemotherapy prediction problem, five cancer types [colon adenocarcinomas (COAD), pancreatic adenocarcinoma (PAAD), bladder carcinoma (BLCA), sarcoma (SARC), and breast invasive carcinoma (BRCA)] totaling 2, 606 tumors. We selected the five cancer types based on two criteria: (i) a sufficient number (at least 65) of paired tumor-transcriptome and clinical data samples available for the cancer type; and (ii) a sufficient number (at least 180) of tumor transcriptome samples available (regardless of the clinical data availability) for the cancer type. We obtained both the RNA-seq (gene-level) total-read-count-normalized log_2_(1 + *C*) read counts and normalized (fragments per kilobase of transcript per million mapped reads, FPKM (Dillies *et al.*, 2013)) expression data for for 60,483 human genes. To focus the machine-learning on the portion of the tumor transcriptome that had the most variation from tumor to tumor, we identified the top 20% most variable genes as measured by the median absolute deviation (MAD) across tumors, of gene expression in terms of FPKM (we used FPKM for this purpose in order to mitigate bias due to read length and tumor-specific depth of sequencing). For deriving feature-sets for XGBoost prediction directly based on transcript abundances or based on VAE- or PCA encoding, the 20% criterion applied to each of the five cancer types yielded a set of 13,584 genes. We computed MAD using the R package `stats` version 3.4.4 (R Core Team, 2013) with default parameters. After the variance-filtering step, we used the log_2_(1 + *C*) of total-count-normalized count values for the top-20% highest-variance genes (that were selected as described above) to obtain (or encode) feature values. We compared the performance—in terms of minimizing the VAE reconstruction loss (see Sec. 5.3)—of different feature scaling methods (no scaling, min-max normalization, and standardization (Kreyszig *et al.*, 2011)) and selected min-max normalization as the method that we used to rescale gene-level count data for input into the VAE.

### 5.2 *t*-distributed stochastic neighbor embedding (t-SNE)

We computed *t*-SNE embedding components of the tumors using the function `sklearn.decomposition.manifold.TSNE` from the python software package `scikit-learn` version 0.19.1 with parameters init = “pca”, perplexity = 20, learning_rate = 300, and n_iter = 400. For plotting the tumor transcriptome *t*-SNE embeddings, we used the R software package `ggplot2` version 3.1.1.

### 5.3 Variational autoencoder (VAE)

An autoencoder is a type of model that combines “encoder” and “decoder” neural networks to learn a low-dimensional continuous data encoding from which the input signal can be approximately reconstructed (Kramer, 1991). A key advantage of an autoencoder is that it is unsupervised, i.e., it can be trained without labeled examples. Unlike classical autoencoders (e.g., sparse or denoising autoencoders), the variational autoencoder (VAE) is a generative probabilistic model which maps an input vector to a latent-space *random variable* (r.v.). Below, we mathematically define the VAE.

Let 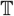 denote the set of tumors for which the VAE is to be fit to the tumor transcriptomes (with 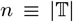) and let *m* denote the number of genes for which transcript abundances are used to represent the tumor transcriptome. After min-max transformation of the tumor transcriptome measurements (Sec. 5.1), each tumor’s transcriptome is represented as a vector ***x*** ∈ [0, 1]^*m*^. Let ***X*** denote the random variable representing the population distribution from which tumor transcriptomes are sampled, and let **X**∈ [0, 1]^*m*×*n*^ represent the composite matrix of all sampled tumor transcriptomes). We aim to learn a VAE that will comprise an encoder and decoder, with the encoder consisting of mean and variance functions ***μ*** : [0, 1]^*m*^ → ℝ^*h*^ and 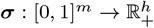, respectively. Together, ***μ*** and ***σ*** map the tumor transcriptome vector ***x***_*t*_ to a *h*-dimensional r.v. ***Z***|***x***_*t*_,

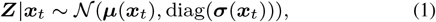

where diag(***m***) is a matrix whose diagonal elements are the elements of the vector ***m***. The decoder is a function ***g*** : ℝ^*h*^ → [0, 1]^*m*^ that, for an outcome ***Z***|***x***_*t*_ = ***z***_*t*_ ∈ ℝ^*h*^, maps

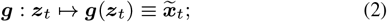

the tilde on 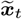 denotes that it is the decoded data for the tumor transcriptome ***x***_*t*_. A good autoencoder should have low reconstruction error *L*, which is convenient to define in terms of the *p*-norm of the difference between the tumor transcriptome data ***x****_t_* and the reconstructed data 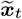, i.e., 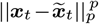 where ‖ ‖_*p*_ denotes the *p*-norm. However, this definition of the reconstruction error is only deterministic in the context of a specific outcome ***Z***|***x***_*t*_ = ***z***_*t*_. Thus, it is conventional to define the reconstruction error as an *expectation value* over outcomes of ***Z***|***x***_*t*_,

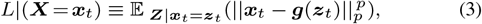

where 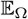 represents an expectation value over a space of outcomes Ω. It should be noted the above representation of the reconstruction error is in terms of the outcome, ***z***_*t*_, of a r.v. (***Z***|***x***_*t*_) whose distributional parameter functions ***μ*** and ***σ*** have hyperparameters (neural network coefficients) that will be fitted. Because Eq. 3 is ill-suited to backpropagation, it is helpful to recast it in terms of a new random variable ***ε***_*t*_ that depends on ***Z***|***x***_*t*_ by

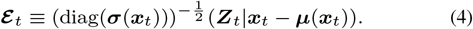

It follows from Eq. 4 and Eq. 1 that ***ε****t* is standard multivariate normal,

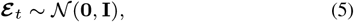

where **I** is the *h* × *h* identity matrix, and thus, ***ε***_*t*_ does not depend on ***μ***, ***σ***, or *t*. We therefore drop the subscript *t* and simply denote the rescaled latent-space random variable as ***ε***. Solving Eq. 4 for ***Z***|***x***_*t*_ and applying it to Eq. 3, the reconstruction error *L*|(***X*** = ***x***_*t*_) can be represented by

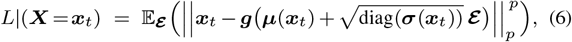

which is amenable to backpropagation because the only r.v. in it is ***ε***, whose distributional parameters do not depend on the neural network coefficients that we will be varying. In practice, rather than computing the multivariate integral over outcomes of ***ε***, *L*|(***X*** = ***x***_*t*_) is typically approximated by averaging over a limited number *J* of samples from ***ε***,

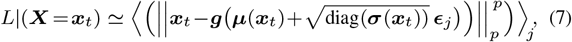

where 〈〉_*j*_ denotes average over *j* ∈ {1, …, *J*} and ***∈***_*j*_ is sample *j* from ***ε***. Following Way and Greene (2017), we used a number of samples that is equivalent to the dimension of the transcriptome, i.e., *J* = *m*. For the case of *p* = 2 (i.e., L2 norm), minimizing *L*|(***X*** = ***x***_*t*_) as defined above is equivalent to maximizing the expectation value of the log-likelihood log(*P* (***g***(***Z***) = ***x***_*t*_ | ***X*** = ***x***_*t*_)). However, following Way and Greene (2017) and consistent with empirical evidence (Sec. 2.3), for our five-cancer study of the utility of a VAE-based approach for response-to-chemotherapy prediction, as well as for the pan-cancer *t*-SNE analysis (Sec. 2.1), we chose to use L1 reconstruction loss, i.e., *p* = 1 in Eq. 3.

The reconstruction loss measures bias error, whose minimization must be balanced against the simultaneous goal of controlling variance error through regularization. In the VAE, regularization requires incentivizing (in the learning of ***μ***, ***σ***, and ***g***) the latent space distributions of ***Z***|***x*** to be close to standard multivariate normal. This is accomplished by assigning a penalty based on the Kullback-Leibler divergence between the distribution of ***Z***|***x***_*t*_ and the target distribution ***ε***, represented by *D*_KL_(*P* (***Z***|***x***_*t*_) || *P* (***ε***)). This regularization is analytically tractable (Duchi, 2007), and for a given tumor *t* yields (see Supplementary Note, Eq. S2) the following regularization function:

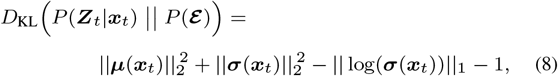

where log(***σ***_*t*_) denotes an element-wise log and ‖ ‖_1_ is the L1 norm.

Fitting the VAE to **X** requires selecting ***μ***, ***σ***, and ***g*** from their respective function spaces; in practice, we search over functions that can be represented using a neural network for ***μ*** and ***σ*** (parameterized by the vector ***θ***)^1^ and a neural network for the function ***g*** (parameterized by the vector ***ϕ***). Exploring the space of functions ***μ***_***θ***_, ***σ***_***θ***_, and ***g***_***ϕ***_ corresponds to computationally searching for the vector pair 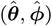 that together minimize the joint (over all tumors) sum of the tumor-specific reconstruction loss and the regularization penalty,

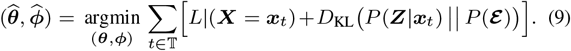

Applying Eqs. 6, 7, and 8, and setting *p* = 1 as discussed above, we obtain the explicit formula for fitting a VAE to **X**,

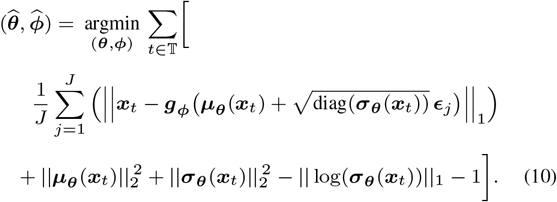

We implemented Eq. 10 in Tensorflow version 1.4.1 with Keras version 2.1.3 as the model-level library. We solved Eq. 10 using the Adam optimization algorithm (Kingma and Ba, 2014) (with batch normalization) from the python package `keras-gpu` version 2.1.3 with parameters learning_rate = 2 × 10^−3^, beta_1 = 0.9, and beta_2 = 0.999, to obtain 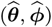. Then, for each tumor *t*, we used a single sample ***Z***|***x***_*t*_ = ***z***_*t*_ from the distribution 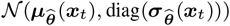 as the final latent-space encoding of the tumor to be used for supervised learning (Sec. 5.6).

### 5.4 Labeling tumors based on response to chemotherapy

From Xena and cBioPortal (Cerami *et al.*, 2012; Gao *et al.*, 2013), we obtained and combined TCGA clinical data (where available) for the patients whose tumor transcriptomes we acquired (see Sec. 5.1). From Xena, we extracted the variables submitter_id.samples, therapy_type, and measure_of_response; from cBioPortal, we extracted the variables Sample_ID, Disease.Free.Status, and Pharmaceutical.Therapy.Indicator. We co-analyzed the Xena- and cBioPortal-obtained clinical data to label tumors “responded” (*y* = 0) or “progressive” (*y* = 1), by assigning *y* = 0 when the clinical record had Complete response or partial response in the measure_of_response column of the clinical data from Xena, or with value DiseaseFree in the Disease.Free.Status column of the clinical data from cBioPortal while therapy type is recorded as Chemotherapy in both. We assigned *y* = 1 to tumors whose clinical records had values Radiographic progressive disease, Clinical progressive disease, or stable disease in the Xena clinical data column measure_of_response, or had value Recurred/ progressed in the cBioPortal data column Disease.Free.Status while the therapy_type is recorded as Chemotherapy in both files. This yielded 806 labeled tumors out of 2,606 total. A total of 39 different drugs were used to treat the 794 patients (see Supplementary Note, Table S1).

### 5.5 VAE model architectures

We trained six transcriptome-encoding VAEs based on four VAE architectures, the pan-cancer VAE architecture (for the 32-cancer unsupervised analysis, see Sec. 2.1) and three cancer type-specific VAE architectures for response-to-chemotherapy prediction (Sec. 2.4) (one of which was used for three different cancer types, BLCA, BRCA, and PAAD, and the others of which were cancer type specific for COAD and SARC). For the pan-cancer VAE, we used a latent space dimension *h* = 50 and three fully connected layers each for the encoder and decoder. For the cancer type-specific VAE architectures, we again used the same number of fully-connected layers in the encoder as in the decoder (Table 3).

**Table 3.** VAE architectures used for predicting chemotherapy response (*h*, latent space dimension; “layers”, # of layers used in the encoder/decoder).

| Name | Cancer types | $h$ | Layers |
| --- | --- | --- | --- |
| VAE-1 | BLCA, BRCA, PAAD | 50 | Six |
| VAE-2 | COAD | 650 | Two |
| VAE-3 | SARC | 500 | Two |

### 5.6 Regularized gradient boosted decision trees (XGBoost)

For predicting whether or not (based on its transcriptome-derived feature-set: raw, PCA, or VAE) a tumor would respond to chemotherapy, we used XGBoost (Chen and Guestrin, 2016), an efficient implementation of regularized gradient boosted decision trees. We used the binary classifier function `XGBClassifier` from the python software package `xgboost` version 0.80, with `gamma=0`. We tuned eight hyper-parameters (Table 4) by exhaustive grid-search with five-fold cross-validation, using `sklearn.model_selection.GridSearchCV` from `scikit-learn` version 0.19.1. To obtain feature importance scores, we used `get_score` with `importance_type = cover`.

**Table 4.**
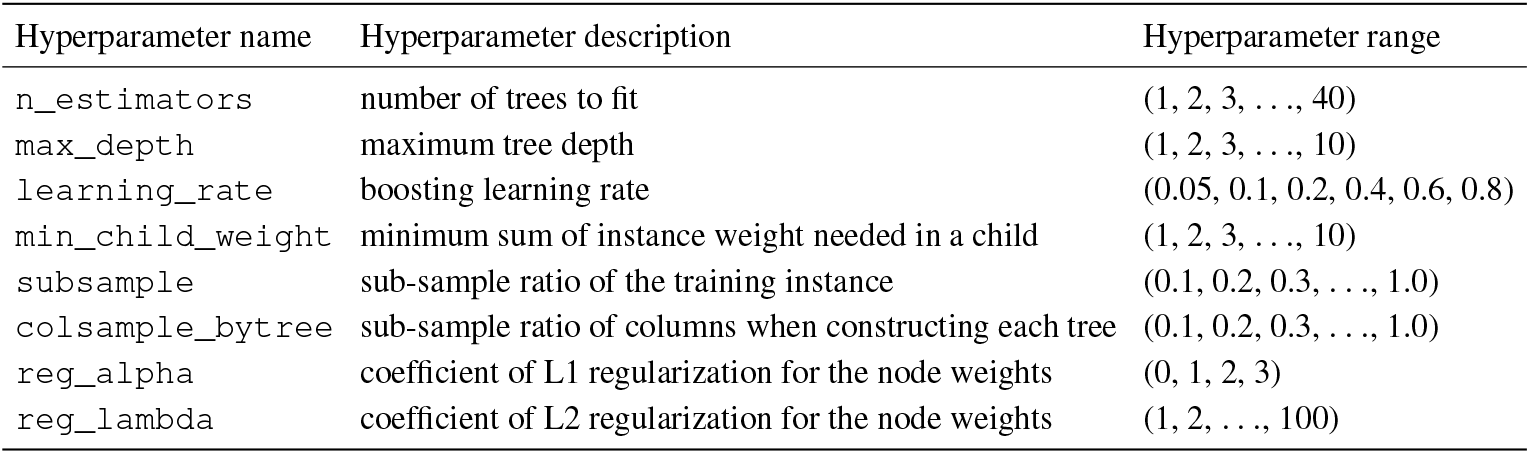
XGBoost classification algorithm hyperparameters and hyperparameter ranges used in grid-search tuning.

### 5.7 Area Under ROC Curve (AUROC)

For computing the AUROC (i.e., sensitivity versus false positive error rate curve), we used the function `metrics.roc_auc_score` from the python software package `scikit-learn` version 0.19.1 with parameter `average=“weighted”`. We logit-transformed AUROC values before testing (using two-tailed Welch’s *t*-test and the Wilcoxon signed rank test)

For the L1 vs. L2 analysis (Fig. 2.3), we carried out 30 replications of five-fold cross-validation; within each replication, across the five folds, we obtained prediction scores for each tumor from the fold in which the tumor was in the test set, enabling us to compute an overall AUROC within each replication. For each training data set, we have done 30 replications of five-fold cross-validation by altering the random seed used for assign split of data during cross-validation. We have conducted the same procedure for five different cancer types (BLCA, BRCA, COAD, PAAD, SARC) as shown in the panel names of Figure 4.

### 5.8 Principal component analysis (PCA)

For PCA, we used the function `decomposition.PCA` (with parameters svd_solver = “full″) and n_components = 0.9 (90% variance, yielding 387 components) from the python package `scikit-learn` version 0.19.1. For plotting, we used `matplotlib` version 2.1.2.

## Funding

SAR acknowledges support from the Animal Cancer Foundation.

## Footnotes

1 Note, functions ***μ*** and ***σ*** are just two different outputs of the encoding neural network, differing only at the final layer, and thus for simplicity of notation we represent them as having a common parameter vector ***θ***.

## Notes

https://github.com/ATHED/VAE_for_chemotherapy_drug_response_prediction

### Competing Interest Statement

The authors have declared no competing interest.

