## Supplementary material for "Predicting chemotherapy response using a variational autoencoder approach"

### Predicting chemotherapy response using a variational autoencoder approach: Supplementary Material

QI WEI AND STEPHEN A. RAMSEY

January 4, 2021

#### A. Supplementary Figure

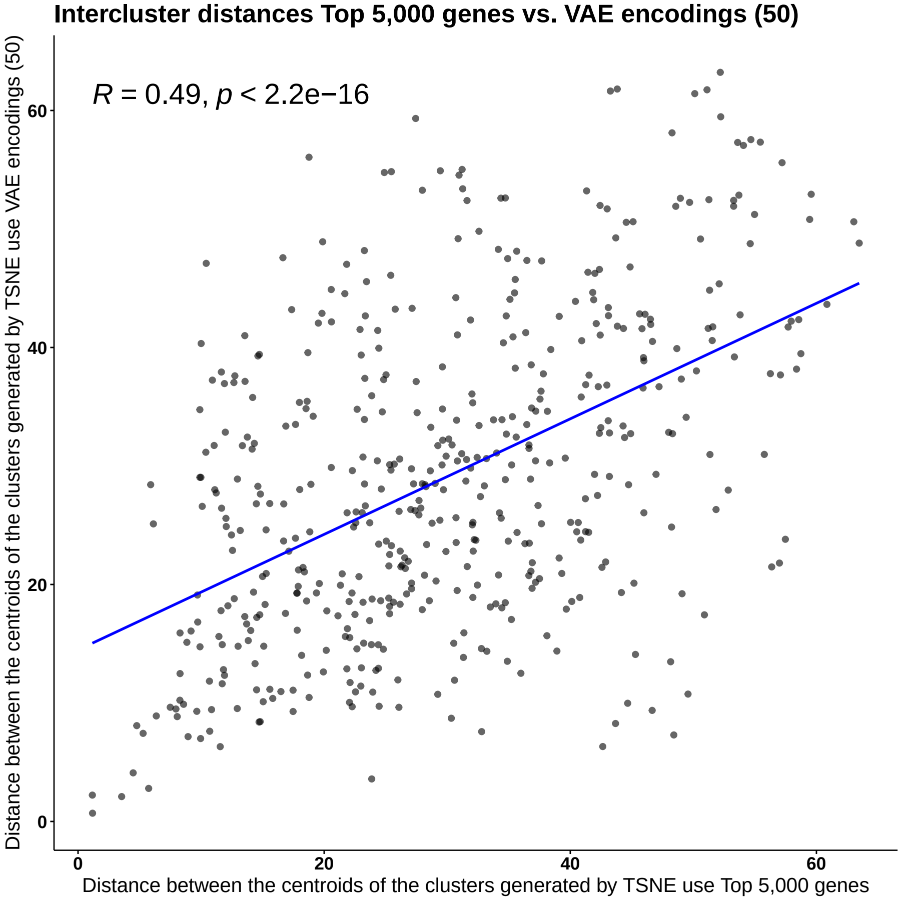


**Fig. S1.** Each mark corresponds to an unordered pair of cancer types (from the 32 cancer types in Fig. 1 in the main article). The horizontal axis measures the distance between the two clusters' centroids based on the *t*-SNE visualization of the tumor expression levels of the top 5,000 genes, as in Fig. 1A (main article). The vertical axis measures the distance between the two clusters' centroids based on the *t*-SNE visualization of the VAE encoding (with latent space dimension $h = 50$) of the expression levels of the top 5,000 genes, as in Fig. 1B (main article).

#### B. Supplementary Table

**Table S1.** Therapeutic agents that were used to treat patients in the TCGA clinical dataset (Hutter and Zenklusen, 2018; Goldman *et al.*, 2019) (see Sec. 5.4), for the five cancer types for which our model was trained to predict response. In some cases, the specific chemotherapeutic agent was not available in the TCGA clinical record.

| cancer type | therapeutic agents |
|---|---|
| breast invasive carcinoma (BRCA) | Goserelin, Trastuzumab, Fluorouracil, Cyclophosphamide, Gemcitabine, Docetaxel, Doxorubicin, Capecitabine, Paclitaxel, Bevacizumab, Methotrexate, Vinorelbine, Etoposide, Carboplatin, Tamoxifen, Pemetrexed, Lapatinib, Everolimus, Epirubicin, Letrozole |
| colon adenocarcinomas (COAD) | Fluorouracil, Oxaliplatin, Irinotecan, Capecitabine, Leucovorin, Cetuximab, Regorafenib |
| bladder carcinoma (BLCA) | Cisplatin, Etoposide, Paclitaxel, Carboplatin, Gemcitabine, Doxorubicin, Fluorouracil, Docetaxel, Methotrexate, Ifosfamide, Vinblastine, Vorinostat, Platinum, Vinorelbine, Vandetanib |
| pancreatic Adenocarcinoma (PAAD) | Gemcitabine, Fluorouracil, Oxaliplatin, Irinotecan, Capecitabine, Cisplatin, Erlotinib, Paclitaxel |
| sarcoma (SARC) | Gemcitabine, Pazopanib, Docetaxel, Temozolomide, Cisplatin, Ifosfamide, Palbociclib, Dacarbazine, Alisertib, Doxorubicin, Carboplatin, Pemetrexed, Sorafenib, Tivozanib |

#### C. Supplementary Note

Using the Kullback-Leibler divergence as the measure of deviation and assuming the latent prior is iid Gaussian, the inputs are two multivariate normal distributions of dimension $h$, for which in general the KL divergence formula (Duchi, 2007) is written as:

$$D_{\mathrm{KL}}(p_1||p_2) = \frac{1}{2}[\log \frac{|\Sigma_2|}{|\Sigma_1|} - h + tr\{\Sigma_2^{-1}\Sigma_1\} + (\mu_2 - \mu_1)^T \Sigma_2^{-1}(\mu_2 - \mu_1)], \tag{S1}$$

where $p_1 = \mathcal{N}(\mu_1, \Sigma_1)$ and $p_2 = \mathcal{N}(\mu_2, \Sigma_2)$. In VAE model, we have $p_1 = P(\boldsymbol{Z}|\boldsymbol{x})$ and $p_2 = P(\boldsymbol{\mathcal{E}})$) and Equation S1 can be written as:

$$\begin{aligned} D_{KL}(P(\boldsymbol{Z}|\boldsymbol{x})||P(\boldsymbol{\mathcal{E}})) &= \frac{1}{2}[\log \frac{|I|}{|\Sigma|} - h + tr\{I^{-1}\Sigma\} + (\vec{0} - \mu)^T I^{-1}(\vec{0} - \mu)] \\ &= \frac{1}{2}[-\log|\Sigma| - h + tr\{\Sigma\} + \mu^T\mu] \\ &= \frac{1}{2}[-\log\prod_i \sigma_i^2 - h + \sum_i \sigma_i^2 + \sum_i \mu_i^2] \\ &= \frac{1}{2}[-\sum_i \log\sigma_i^2 - h + \sum_i \sigma_i^2 + \sum_i \mu_i^2]. \end{aligned} \tag{S2}$$
